# Model-Guided Engineering of NS1-Truncated Influenza OP7 Particles with Enhanced Interferon-Mediated Antiviral Activity

**DOI:** 10.64898/2026.09.18.752587

**Authors:** Julita Piasecka, Daniel Rüdiger, Udo Reichl, Sascha Young Kupke

## Abstract

Defective interfering particles (DIPs) represent promising antivirals against influenza A virus (IAV), yet strategies to further enhance their antiviral activity remain largely unexplored. Here, we engineered a next-generation “OP7” DIP by truncating the viral interferon (IFN) antagonist non-structural protein 1 (NS1) to enhance IFN-mediated antiviral activity. A previously validated mathematical multiscale model predicted that loss of NS1 function would enhance antiviral activity, providing a mechanistic rationale for this design. We engineered “OP7-trNS1” DIP, established a cell culture-based production process yielding infectious virus-free material, and evaluated its antiviral activity in human lung epithelial cells *in vitro*. OP7-trNS1 inhibited IAV replication by approx. two orders of magnitude more effectively than the parental OP7 DIP while inducing a stronger IFN response, consistent with the model predictions. Together, these findings demonstrate that reducing viral IFN antagonism is a viable strategy for rationally enhancing DIP antiviral activity and establish OP7-trNS1 as a promising candidate for further preclinical development.

## INTRODUCTION

In the era of globalization, respiratory viral infections such as those caused by the influenza A virus (IAV) can spread rapidly, leading to large-scale outbreaks and annual epidemics. Occasionally, a global pandemic occurs with even higher number of infections, some of which progress to severe disease or fatal outcomes [1, 2]. Rapid genetic adaptations of IAV pose a major challenge to current standards of care. For instance, antigenic drift driven by mutations in surface glycoproteins lead to vaccine antigen mismatches over time [1]. Therefore, vaccine compositions require regular, seasonal updates. However, the adaptive immunity requires two to four weeks to unfold full efficacy. Therefore, due to their fast mode of action, small-molecule antivirals are additionally used to complement vaccination and to increase pandemic preparedness. However, viral evolution facilitates also the emergence of antiviral resistance through alterations in drug target sites. Therefore, novel antiviral strategies are under scrutiny to address unmet needs of the healthcare system. In this context, virus-derived defective interfering particles (DIPs), posing a high barrier to the evolution of viral resistance [3–6], have attracted attention due to their ability to suppress infections. For IAV, several *in vivo* studies demonstrated the antiviral potential of IAV DIPs [7–9].

Conventional DIPs derived from IAV harbor a large internal deletion within one of their eight viral RNA (vRNA) genome segments [4]. OP7 is a nonconventional type of IAV DIP, previously identified in our laboratory, characterized by the presence of a hypermutated genomic segment 7 (S7) vRNA [10]. Both types of genomic alterations render DIPs replication incompetent, likely due to the loss of essential protein functions. However, during coinfection with infectious IAV, DIPs show enhanced replication of the defective interfering (DI) vRNA. This impedes the replication of infectious homologous viruses through competition for cellular and viral resources, a process termed replication interference [11–14]. Simultaneously, DIPs activate interferon (IFN) responses in host cells [15–17], which contribute to the antiviral efficacy of DIPs [18–23]. The IFN-dependent antiviral effect has also been demonstrated to inhibit nonhomologous viruses [24–26]. For instance, IAV DIPs can suppress the replication of other respiratory viruses, including that of the influenza B virus, respiratory syncytial virus (RSV), and SARS-CoV-2 [27–29]. This broad-spectrum antiviral activity may be particularly valuable during pandemics, before vaccines or virus-specific antivirals become available.

In previous studies, OP7 has demonstrated enhanced antiviral potential compared to conventional IAV DIPs [27, 30, 31]. In addition, a modified reverse genetics workflow has been established for the reconstitution of an OP7 chimera DIP that harbored both, a deletion in segment 1 (S1) vRNA and the hypermutated segment 7 of OP7 (S7-OP7) vRNA [7, 32]. Subsequently, a high-yield cell culture-based production process for the OP7 chimera DIP was developed in laboratory-scale bioreactors [32]. Importantly, the production of an OP7 chimera preparation free of infectious viruses was possible, a significant advantage for safe administration and potential regulatory approval. The produced OP7 chimera DIP material was tested in murine studies, which proved high *in vivo* tolerability and antiviral efficacy [7].

In addition, considerable research efforts have focused on attenuated IAV mutants containing modifications of the viral non-structural protein 1 (NS1), an IFN antagonist encoded by the genomic segment 8 (S8) of IAV. NS1 performs multiple functions during the viral replication cycle, most notably antagonizing the type I IFN response by impairing vRNA recognition, preventing downstream IFN signaling, and disturbing cellular mRNA processes [33, 34]. As a result, NS1 mutant viruses induce high levels of IFN locally upon infection, which inhibits viral replication and leads to attenuated viral growth. NS1-modified viruses are also potent activators of antigen-presenting cell function, which make them even more immunogenic [35, 36]. Therefore, they represent promising candidates for new live attenuated influenza vaccine (LAIV) designs (reviewed in [37]).

In this study, we aimed to enhance the antiviral activity of OP7 by reducing its ability to antagonize the host IFN response. We therefore designed OP7-trNS1, a chimeric DIP combining the hypermutated S7-OP7 vRNA and a truncated S1 vRNA with a S8 vRNA sequence variant encoding a truncated NS1 protein (NS124) in a single viral particle (OP7-trNS1). A previously validated multiscale model predicted that loss of NS1 would enhance IFN-mediated antiviral activity, providing a mechanistic rationale for this design. We then successfully reconstituted OP7-trNS1, developed a cell culture-based production of OP7-trNS1 and assessed its antiviral activity and IFN induction in human lung epithelial cells *in vitro*. OP7-trNS1 showed substantially enhanced antiviral activity compared with parental OP7, consistent with the model prediction. Together, this study demonstrates a model-guided strategy for rationally enhancing the antiviral activity of DIPs by reducing viral antagonism of host innate immunity.

## RESULTS

### Model simulations predict enhanced antiviral activity of NS1-deficient OP7

To test whether loss of NS1-mediated IFN antagonism could enhance OP7 antiviral activity, we used our previously validated mathematical multiscale model of infectious standard virus (STV) and OP7 coinfection [38]. This multiscale model describes the coinfection, replication and release of STV and OP7 particles at the intracellular and cell population level (Figure 1a). Furthermore, it considers the antiviral activity of the IFN system as well as the immunosuppressant effect of NS1. In short, pattern recognition receptors (PRRs) detect incoming viral ribonucleoproteins (vRNPs), harboring the genomic vRNAs, and initiate a signaling cascade that leads to the synthesis and secretion of IFN-β. IFN-β then activates IFN-α/β receptors (IFNAR) that induce the expression of antiviral IFN-stimulated genes (ISGs) via the Janus kinase, signal transducer and activator of transcription proteins (JAK/STAT) pathway, including human myxovirus resistance protein A (MxA). MxA is a key ISG-induced effector protein limiting IAV infection by inhibiting the nuclear import of vRNPs [38–40]. The viral NS1 protein can bind to PRRs to inhibit the activation of the IFN system, preventing its antiviral activity. The model was calibrated to *in vitro* experimental data derived from human lung epithelial Calu-3 cells [41].

**Figure 1.**
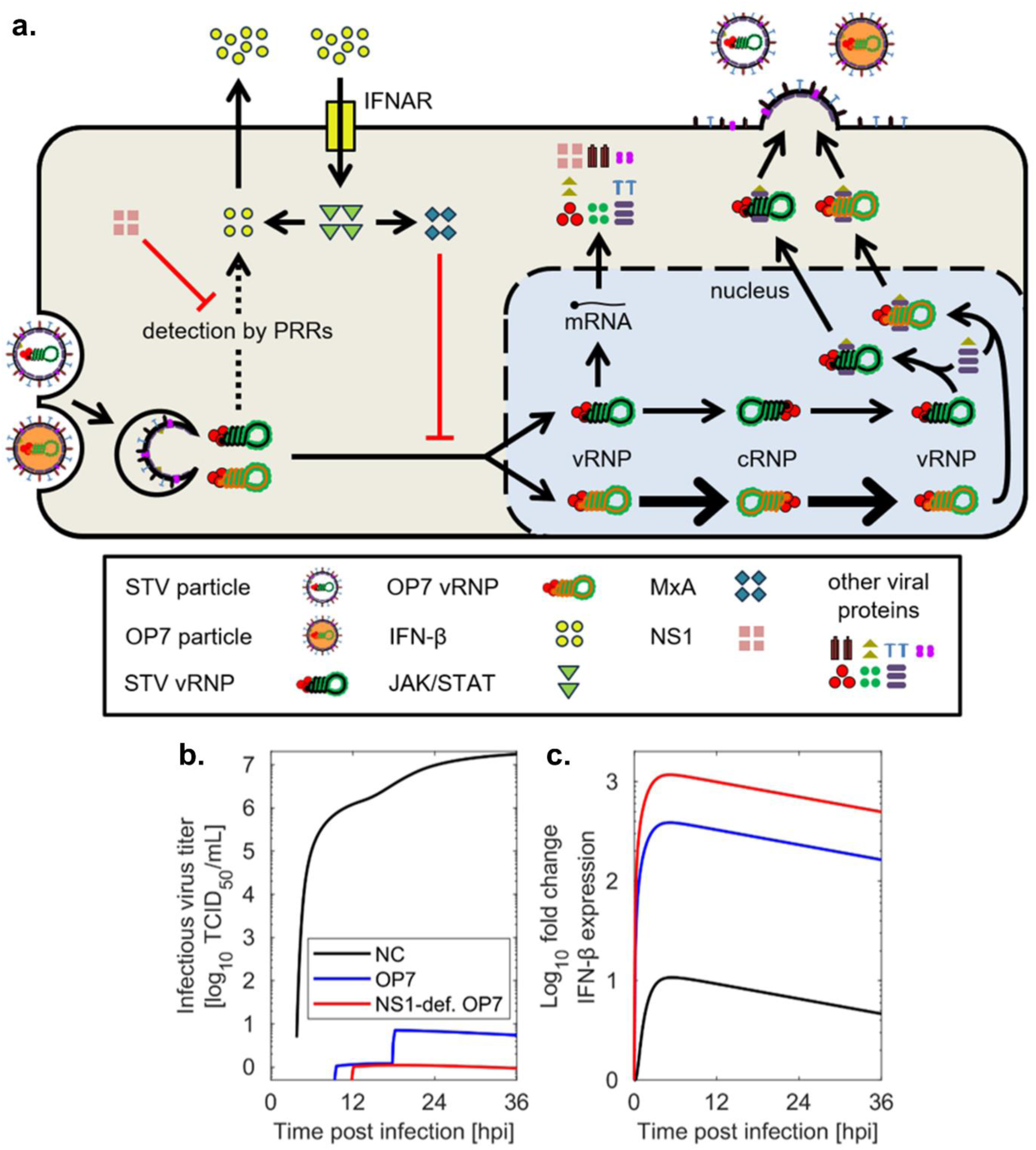
STV and OP7 coinfection model and predicted impact of NS1 loss on antiviral activity. (a) The intracellular model describes virus entry, viral replication and protein synthesis, the IFN response, and progeny virion release of STV-infected and OP7 coinfected cells. Furthermore, the model simulates viral spread across a Calu-3 cell population (not shown). Figure adapted from [38]. (b) Simulated infectious virus titers for (co)infections at STV MOI of 0.2 without OP7 (NC), and with high doses (500 OP7 particles per cell) of regular OP7 or NS1-deficient OP7. (c) Simulated IFN levels for (co)infections at STV MOI of 0.2 without OP7, with regular OP7 or NS1-deficient OP7.

To estimate the impact of removing NS1 expression from OP7 particles, we simulated coinfections of STV and OP7 as well as an OP7 unable to express NS1 (referred to as NS1-deficient OP7). In our simulations, coinfections with NS1-deficient OP7 resulted in lower infectious progeny virus release than coinfections with OP7, indicating an increased antiviral activity of NS1-deficient OP7 (Figure 1b). These differences were dependent on the multiplicity of infection (MOI), with lower STV MOIs leading to larger differences in inhibition by NS1-deficient OP7 compared to OP7 (Supplementary Figure S1).

In Figure 1b, we show simulations of an STV infection at an MOI of 0.2 and coinfections with OP7 or NS1-deficient OP7. The model predicts that coinfection with OP7 reduces infectious virus release (indicated by the 50% tissue culture infectious dose (TCID_50_) titers) by 6 logs while coinfection with NS1-deficient OP7 reduces titers by 7 logs. Further, the model predicts higher IFN-β levels when NS1-deficient OP7 is used for coinfection compared to the regular OP7 (Figure 1c), because the IFN antagonist NS1 is not expressed. This induces an increased production of MxA, inhibiting nuclear import of STV vRNA and subsequent virus replication, which explain the enhanced titer reductions. This effect is more pronounced at lower MOIs (Supplementary Figure S1), which would be especially beneficial for physiologically relevant coinfection conditions as low MOIs mirror natural human infections [42].

Taken together, model simulations predict that loss of NS1 function would enhance OP7 antiviral activity by strengthening the IFN-mediated antiviral state.

### Rational design and successful reconstitution of the NS1-truncated OP7 (“OP7-trNS1”)

We next aimed to generate an OP7 DIP unable to express the full-length NS1 using plasmid-based reverse genetics. For this, we used a system for reconstitution of conventional IAV DIPs free of infectious viruses [43] and a modified version for reconstitution of OP7 chimera DIPs [7]. Recognizing the challenges associated with the reverse genetics-based virus reconstitution, we generated new constructs stepwise, introducing one additional genomic change at a time to ensure its compatibility with virus replication or to allow further optimization of the conditions. Figure 2a depicts all IAV genomic alterations considered in this study.

**Figure 2.**
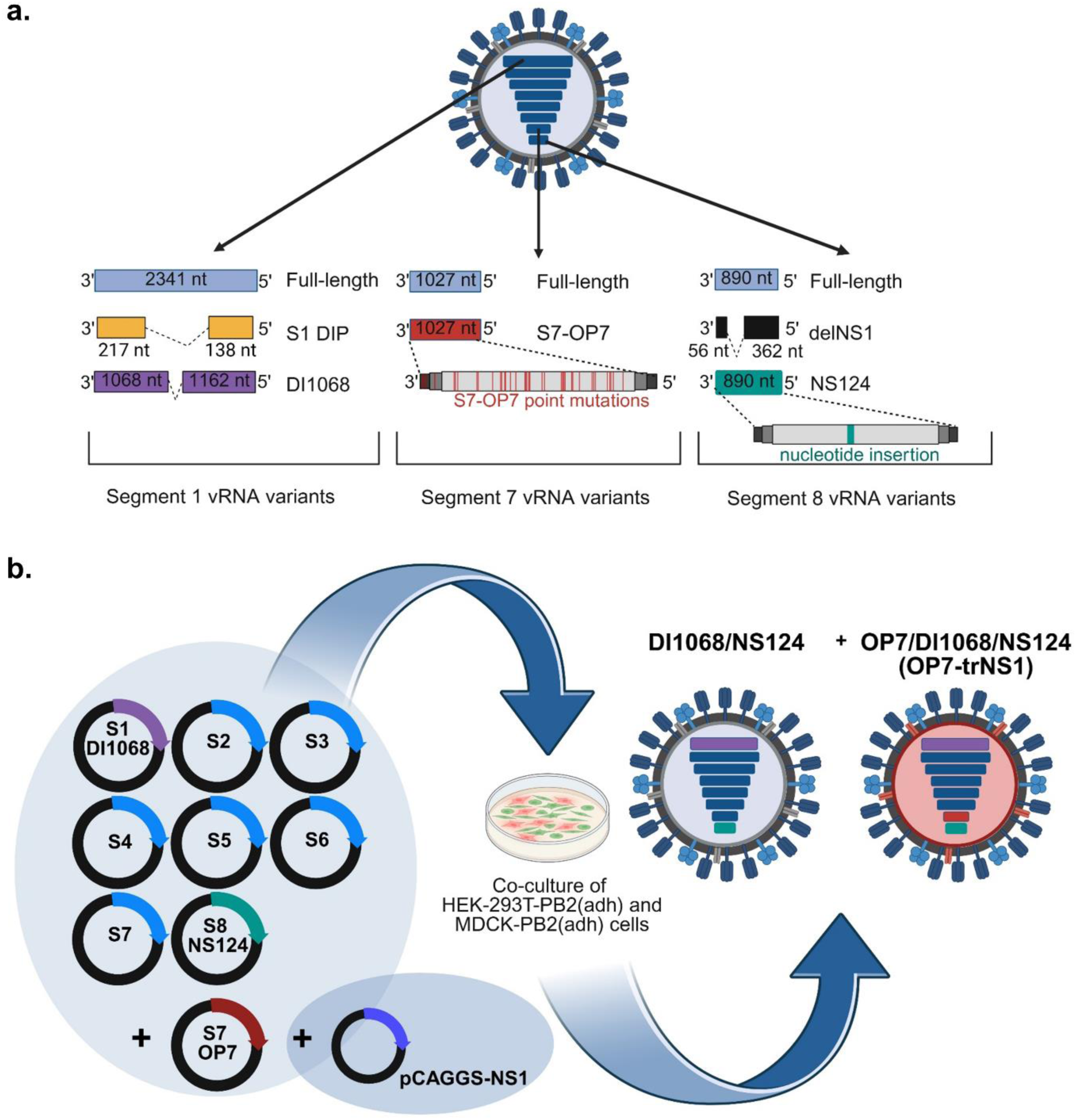
Design and plasmid-based reconstitution of OP7-trNS1. (a) Structure of WT vRNA segments and all segment variants, which were tested in rescue experiments in this study (Table 1). (b) Rescue of OP7/DI1068/NS124 (OP7-trNS1). Nine plasmids provided a deleted S1 vRNA (DI1068), S7-OP7 vRNA, a mutated S8 vRNA (NS124) and S2–7 WT vRNAs. An additional plasmid (pCAGGS-NS1) provided the NS1 protein expression *in trans*, required during rescue in order to allow for sufficient virus replication. Co-transfection was performed in a co-culture of complementing, PB2-expressing cells (HEK-293T-PB2(adh) and MDCK-PB2(adh)), as DI1068 vRNA contains an internal deletion in S1 vRNA making it unable to express the full-length, functional PB2 protein. As a result, two types of DIPs were reconstituted in one virus population: DI1068/NS124 and OP7/DI1068/NS124 (OP7-trNS1) DIPs. Here, the defective S7-OP7 vRNA in OP7-trNS1 is complemented by the fully functional S7 WT vRNA from DI1068/NS124, during rescue to enable virus replication, and in coinfections for further propagation in PB2-expressing cells (Figure 4). Figure was created with BioRender.com.

**Table 1.** Virus reconstitution attempts of individual constructs. Genetic composition of the constructs and reconstitution outcome.

| Construct name | Mutated segments | Reconstitution |
| --- | --- | --- |
| S1 DIP | S1 | + |
| DI1068 | S1 | + |
| S1 DIP/OP7 | S1, S7 | + |
| DI1068/OP7 | S1, S7 | + |
| delNS1 | S8 | + |
| NS124 | S8 | + |
| S1 DIP/delNS1 | S1, S8 | - |
| S1 DIP/NS124 | S1, S8 | - |
| DI1068/delNS1 | S1, S8 | - |
| DI1068/NS124 | S1, S8 | + |
| OP7/DI1068/NS124<br>(OP7-trNS1) | S1, S7, S8 | + |

In general, in order to reconstitute conventional DIPs harboring a deletion in S1 vRNA, a co-culture of adherent cells (adh) expressing polymerase basic protein 2 (PB2) was used for the plasmid transfection. DIPs containing a deletion in S1 vRNA are unable to express functional PB2 protein encoded on S1 vRNA. Therefore, the STV-free reconstitution requires complementing, PB2-expressing cells to enable rescue. A successful reconstitution of such DIPs (S1 DIP and DI1068) (Figure 2a and Table 1), was carried out using eight-plasmid transfections. The transfection mixtures contained one plasmid coding for the deleted S1 vRNA and seven encoding wild-type (WT) segments 2-8 (Table 1). S1 DIP contains a large internal deletion in S1 vRNA, previously identified as “Seg 1 gain” [44]. DI1068 contains a relatively short internal deletion in S1 vRNA (<100nt) [45].

Next, we rescued double mutants: S1 DIP/OP7 and DI1068/OP7 (Table 1). For these chimeric DIPs, an additional plasmid encoding the mutated S7-OP7 vRNA was required. This was necessary because the S7 WT vRNA expressed by corresponding plasmid can complement the defect of S7-OP7 vRNA, enabling virus replication and rescue. This nine-plasmid system resulted in the rescue of two types of DIPs in one virus population: conventional DIPs harboring S7 WT vRNA and a deletion in S1 vRNA, and OP7 chimera DIPs, containing S7-OP7 vRNA, a truncated S1 vRNA (S1 DIP or DI1068), and the remaining six WT vRNAs.

Viruses carrying deletions in S8 vRNA are known to be highly attenuated, due to the lack of NS1 protein expression and its abrogated regulatory functions. In particular, the missing IFN antagonism leads to an enhanced antiviral state that hinders viral replication. To overcome these challenges in the abovementioned reverse genetic system, an additional plasmid (pCAGGS-NS1) was included to provide NS1 protein expression *in trans* to facilitate the reconstitution process. We also introduced an A14U mutation in S7 vRNA [46], which was previously described to support replication of influenza viruses lacking NS1. This way, we established an efficient system for the reconstitution of delNS1 and NS124 viruses (Figure 2a, Table 1). DelNS1 is a deletion mutant that cannot express the entire NS1 protein [47]. NS124 contains a single-nucleotide insertion in S8 vRNA, which generates a stop codon [48]. As a result, NS124 expresses a truncated and not fully functional NS1 protein.

Next, we attempted to rescue DIPs with mutated NS1: S1 DIP/delNS1, S1 DIP/NS124, DI1068/delNS1, and DI1068/NS124 (Table 1). From all the tested genomic compositions combining alterations in S1 and S8 vRNAs, only DI1068/NS124 yielded detectable titers following reconstitution (Supplementary Figure S2 and Table 1). This viral mutant contained minimal sequence alterations in both segments, i.e., a short deletion in S1 vRNA and single-nucleotide insertion in S8 vRNA.

Accordingly, a total number of ten plasmids were used for the reconstitution OP7/DI1068/NS124 (Figure 2b). Plasmids encoding for the mutated S7-OP7 vRNA, S7 vRNA with the A14U substitution, S1 vRNA DI1068, S8 vRNA NS124 and the remaining five WT segments were used along with the pCAGGS-NS1 plasmid. This resulted in the rescue of two types of DIPs in one virus population: one containing DI1068 and NS124 (DI1068/NS124), and another containing S7-OP7, DI1068, and NS124 (OP7/DI1068/NS124). The latter DIP is hereafter referred to as OP7 chimera DIPs expressing a truncated NS1 (OP7-trNS1).

In summary, genome constellations containing large internal deletions in both S1 and S8 vRNAs could not be rescued. We therefore retained the S8 vRNA sequence while introducing a stop codon to prevent full-length NS1 expression and reduced the length of the S1 vRNA deletion. This modified genome constellation enabled successful rescue of OP7-trNS1.

### NS1 truncation enhances the antiviral activity of DI1068/NS124 in IFN-competent cells

Before evaluating the more complex OP7-trNS1 construct, we first tested whether the NS1 truncation enhances DIP antiviral activity for the conventional DIP DI1068. OP7-trNS1 represents the most complex DIP construct successfully reconstituted in this study. However, cell culture-based production of similar OP7 chimera DIPs in shake flasks required multiple optimization steps to achieve high yields and enrichment of S7-OP7-containing DIPs in previous studies [7, 32]. Therefore, we initially chose to validate the impact of the NS1 truncation in a simplified and less labor-intensive setup, specifically using DI1068/NS124.

Therefore, we used the rescued material of the DI1068 and DI1068/NS124 constructs and compared their antiviral potential in an interfering assay. In brief, cells were either infected with IAV (strain A/PR/8/34 (H1N1), PR8) alone at an MOI of 0.01 or coinfected with DI1068 or DI1068/NS124 DIPs. In this assay, the inhibition of infectious virus release is an indicator of the antiviral effect triggered by DIP coinfection. MDCK and Calu-3 cells were used to distinguish the effects induced in the absence of a fully functional IFN system against human IAV strains [49] and IFN-competent cells, respectively. In MDCK cells, DI1068 and DI1068/NS124 showed comparable levels of interference with virus replication, reducing the infectious virus release by approx. three orders of magnitude (Figure 3a). In contrast, in Calu-3 cells, DI1068/NS124 exhibited an approx. one order of magnitude greater antiviral effect than DI1068 (Figure 3b). Here, the stronger antiviral effect, which was not present in partially IFN-deficient MDCK cells, appears to be due to diminished IFN antagonism of the DI1068/NS124 DIP.

**Figure 3.**
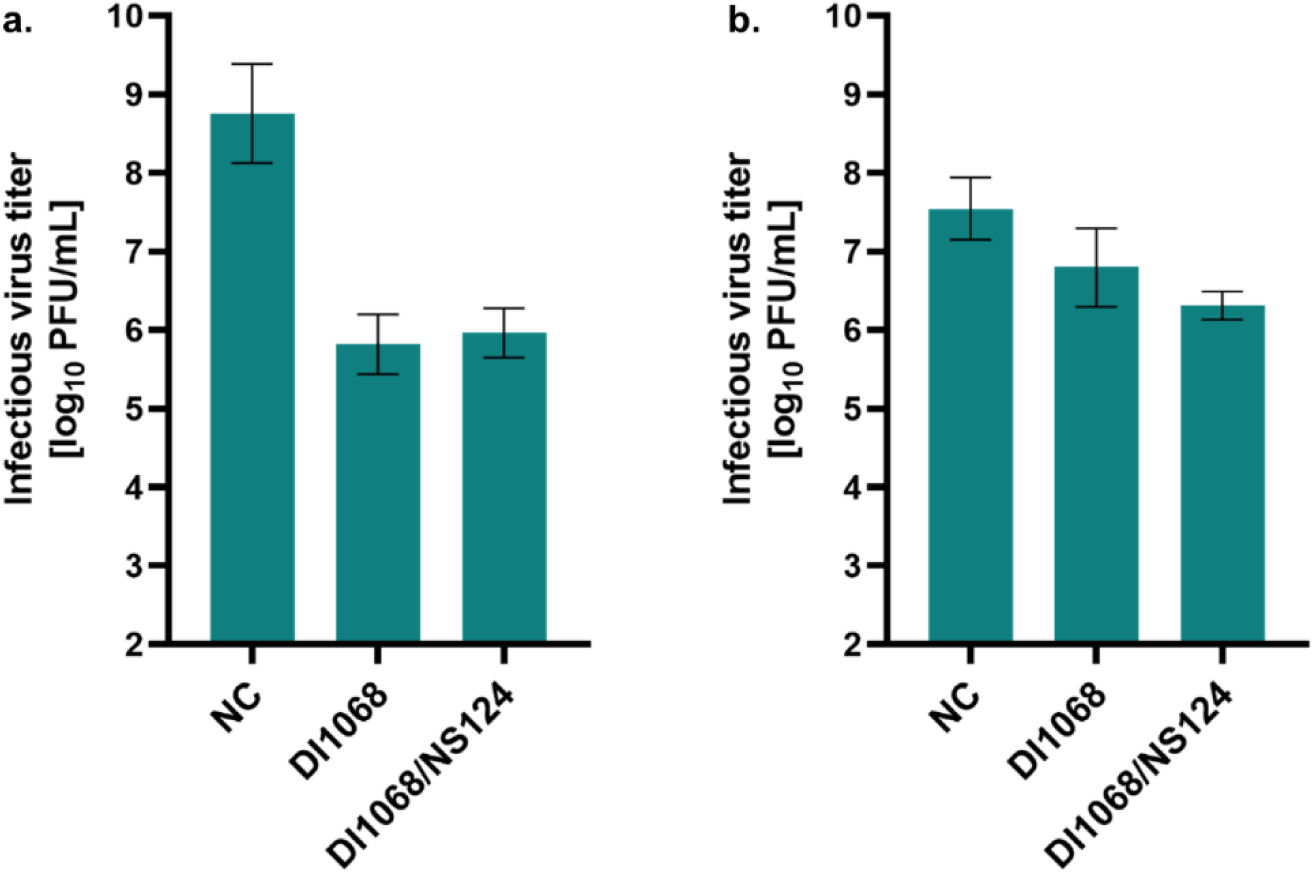
NS1 truncation enhances DI1068 DIP antiviral activity in IFN-competent human lung cells. *In vitro* interference assay was performed in (a) partially IFN-deficient MDCK and (b) IFN-competent Calu-3 cells. Cells were infected with IAV PR8 alone at a MOI of 0.01 (NC) or coinfected with DIPs: DI1068 or DI1068/NS124. Infectious virus release was measured by the plaque assay at 24 hpi. DIP input was normalized and calculated based on the HA titer. Error bars indicate the standard deviation of three independent experiments.

In summary, the NS1 truncation enhanced DI1068/NS124 DIP antiviral activity specifically in IFN-competent Calu-3 cells, consistent with the model prediction that reduced NS1-mediated IFN antagonism enhances DIP antiviral activity (Figure 1). The absence of this effect in partially IFN-deficient MDCK cells further supports an IFN-dependent mechanism.

### Optimization of cell culture-based production enabled high yields and enriched OP7-trNS1 preparations

Next, we established a cell culture-based production process for OP7-trNS1 and optimized the production conditions. The OP7-trNS1 seed virus also contains conventional DIPs (DI1068/NS124) (Figure 2b). In order to enrich OP7-trNS1 during production while maintaining high virus titers, we tested different production MOIs. Similarly, previous OP7 chimera DIP production has been reported to be strongly dependent on the MOI with respect to virus titers and OP7 chimera DIP fractions. We produced OP7-trNS1 DIPs using MDCK-PB2 suspension (sus) cells in shake flasks as previously described [7, 32]. For DI1068/OP7 production, an MOI screening was conducted as well, in order to produce it as a reference for later comparisons (Figure 7).

A range of MOIs from 1E-1 to 1E-6 was tested (Figure 4). For OP7-trNS1 production at MOI 1E-1, the viable cell concentration (VCC) dropped quickly after 16 hours postinfection (hpi) (Figure 4a). For MOIs ranging from 1E-2 to 1E-5, VCC steadily increased reaching a maximum of 3.5 x 10^6^ cells/mL for production at MOI 1E-5. Interestingly, total virus yield, expressed as the hemagglutination (HA) titer, reached similar maximum values of approx. 2.5 for nearly all production MOIs (Figure 4b), except for MOI 1E-6, for which no detectable HA titer was observed. Next, OP7-trNS1 DIP fractions were calculated based on the S7-OP7 and S7 WT vRNA levels in progeny virions, quantified by reverse transcription real-time PCR (RT-qPCR). Analysis revealed the highest OP7-trNS1 fraction (>96%) at production MOIs of 1E-1 and 1E-2 (Figure 4c).

**Figure 4.**
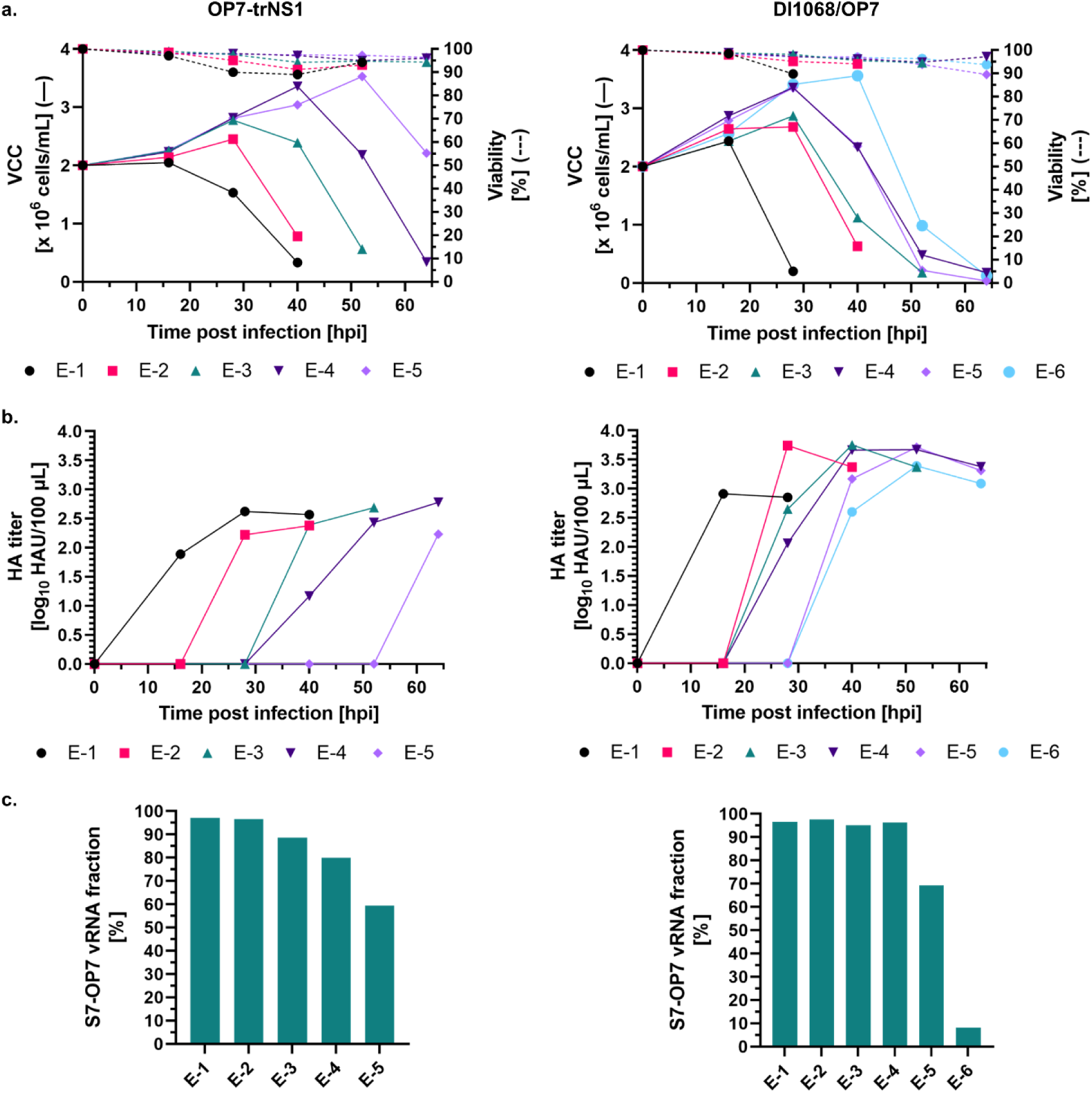
Optimization of cell culture-based production of OP7-trNS1 in shake flasks. MDCK-PB2(sus) cells cultivated in 125 mL shake flasks (50 mL working volume) were infected at MOIs ranging from 1E-1 to 1E-6 after a complete medium exchange. (a) VCC and viability. (b) HA titer. (c) Fraction of OP7-trNS1 and DI1068/OP7 DIPs (calculated based on S7-OP7 and S7 WT vRNA levels of progeny virions, quantified by RT-qPCR). Only selected harvest time points were analyzed (OP7-trNS1: MOI 1E-1/28 hpi, 1E-2/28 hpi, 1E-3/40 hpi, 1E-4/52 hpi, 1E-5/64 hpi; DI1068/OP7: MOI 1E-1/16 hpi, 1E-2/28 hpi, 1E-3/40 hpi, 1E-4/40 hpi, 1E-5/40 hpi, 1E-6/40 hpi).

For the DI1068/OP7 production at MOI of 1E-1, VCC decreased similarly to OP7-trNS1 production after 16 hpi (Figure 4a). At MOIs of 1E-2 and 1E-3, the maximum VCC did not exceed 2.5 x 10^6^ cells/mL, whereas at MOIs of 1E-4, 1E-5, and 1E-6, VCC peaked at approx.

3.5 x 10^6^ cells/mL at 28, 28 and 40 hpi, respectively. Maximum HA titers reached approx. 3.5 for most conditions, except for MOIs of 1E-1 and 1E-6, for which the HA titers were lower (Figure 4b). The highest DI1068/OP7 fractions (>95%) were detected at MOIs between 1E-1 and 1E-4 (Figure 4c). Samples harvested at time points showing high HA titers and minimal cell death (indicated by a decrease in VCC), which alleviates subsequent downstream purification, were selected for the interference assay to evaluate their ability to inhibit virus replication (Figure 5).

**Figure 5.**
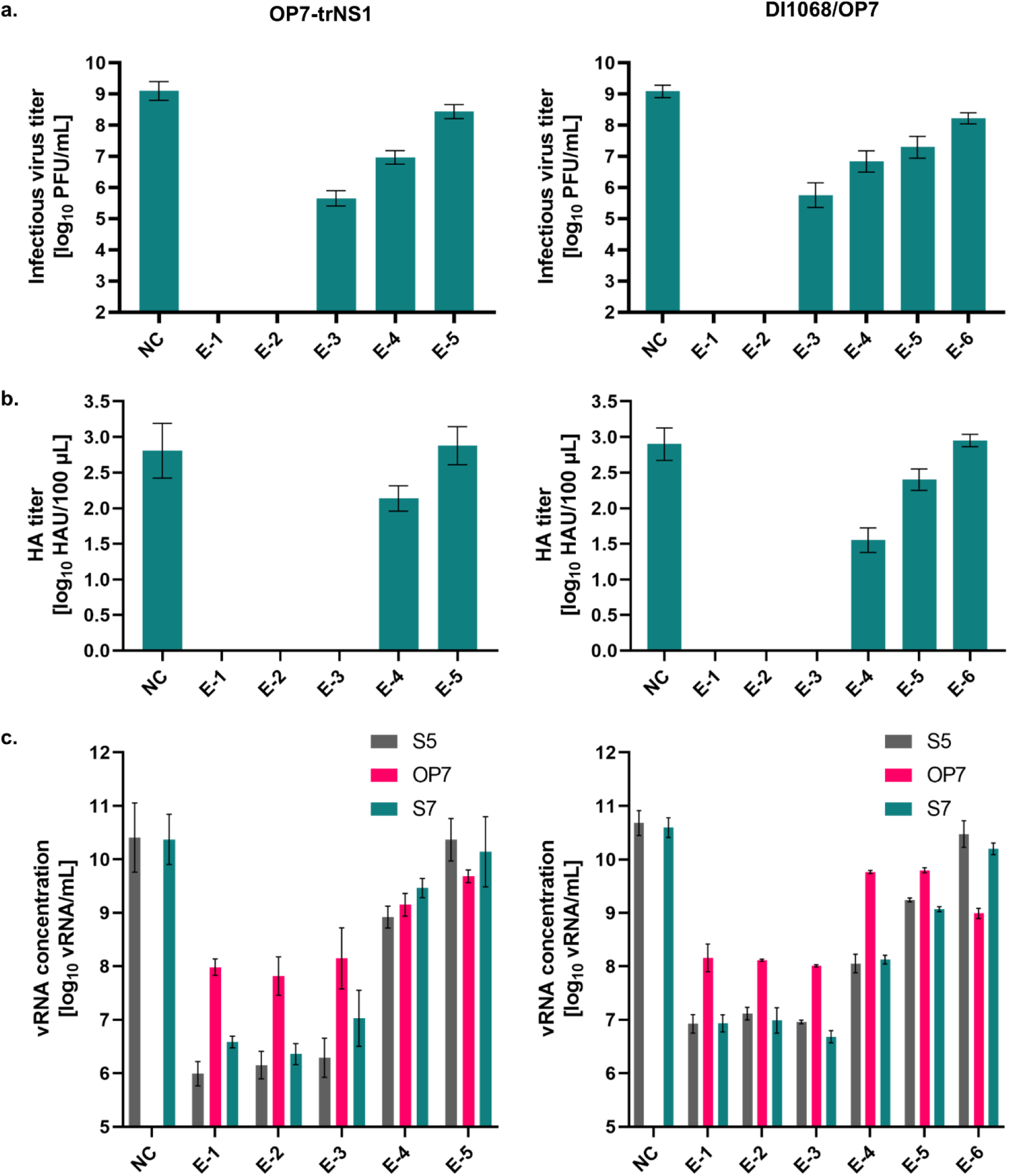
Production MOI-dependent interfering efficacy of OP7-trNS1. An interference assay was performed in MDCK cells. Cells were infected with IAV PR8 alone at a MOI of 0.01 (NC) or coinfected with DIPs (OP7-trNS1 or DI1068/OP7). (a) Infectious virus release measured by plaque assay at 24 hpi. (b) HA titers. (c) vRNA levels in progeny virions in cell culture supernatants, quantified by RT-qPCR. Only supernatants from selected harvest time points were subjected to the interference assay (OP7-trNS1: MOI 1E-1/28 hpi, 1E-2/28 hpi, 1E-3/40 hpi, 1E-4/52 hpi, 1E-5/64 hpi; DI1068/OP7: MOI 1E-1/16 hpi, 1E-2/28 hpi, 1E-3/40 hpi, 1E-4/40 hpi, 1E-5/40 hpi, 1E-6/40 hpi). DIP input was normalized and calculated based on the HA titer. Error bars indicate the standard deviation of three independent experiments.

Coinfection experiments were conducted in MDCK cells to determine the optimal MOI (with respect to inhibition of infectious virus replication) under conditions with minimal influence of IFN responses, thereby focusing on replication interference exerted by S7-OP7 vRNA. Stronger interfering efficacies were observed for the DIP material produced towards higher MOIs (Figure 5). Complete abrogation of IAV PR8 replication was observed for OP7-trNS1 and DI1068/OP7 material produced at MOIs 1E-1 and 1E-2, as reflected by the absence of infectious progeny virus titers (Figure 5a). Toward lower MOIs, the inhibitory effect was gradually becoming less pronounced. Similar trends were observed in the analysis of total virus titers (Figure 5b). Here, no titer was detected up to an MOI of 1E-3, likely due to the higher limit of detection of the HA assay compared to the plaque assay. Further analysis of the vRNA content in progeny virions showed overproportional levels of S7-OP7 vRNA compared to S5 and S7 WT vRNAs in several samples, in accordance with the phenotype typically observed upon OP7 coinfection [10, 31]. The overproportional accumulation was observed across a range of production MOIs of 1E-1, 1E-2, and 1E-3 for OP7-trNS1 and of 1E-1 to 1E-5 for DI1068/OP7.

Considering interfering efficacies, HA titers, times of cell death, and S7-OP7 vRNA fractions, we selected the optimal production MOIs of 1E-1/28 hpi for OP7-trNS1 and 1E-2/28 hpi for DI1068/OP7. For subsequent *in vitro* experiments (Figure 7), both DIPs were produced at optimal production parameters and then purified and concentrated using steric exclusion chromatography (SXC) [50]. Segment-specific RT-PCR analysis confirmed that the produced material showed no significant contamination with other conventional DIPs. Apart from the presence of the truncated S1 vRNA DI1068, no major accumulation of other conventional DI vRNAs in segments S2–S8 was observed (Figure 6) for both productions of OP7-trNS1 and DI1068/OP7. The identity of all genome segments was confirmed by sequencing.

**Figure 6.**
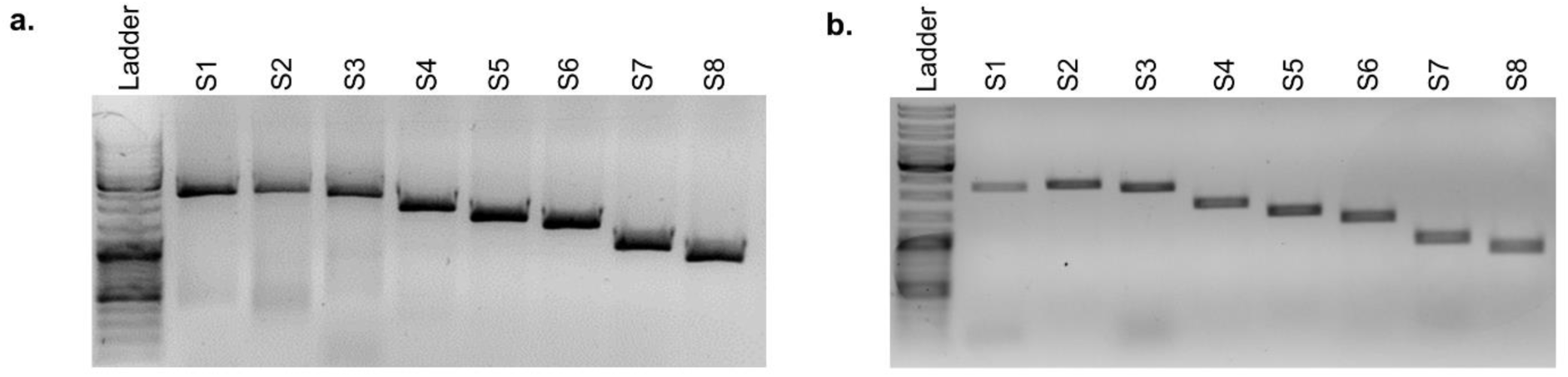
Depletion of contaminating DIPs in the produced OP7-trNS1 material. DIPs were produced at different MOIs in shake flasks (Figure 5). Samples from 28 hpi were subjected to segment-specific RT-PCR and gel electrophoresis. (a) DI1068/OP7 produced at MOI 1E-2. (b) OP7-trNS1 produced at MOI 1E-1. Upper thicker band of the ladder: 3000 bp, middle thicker band: 1000 bp, lower thicker band: 500 bp.

**Figure 7.**
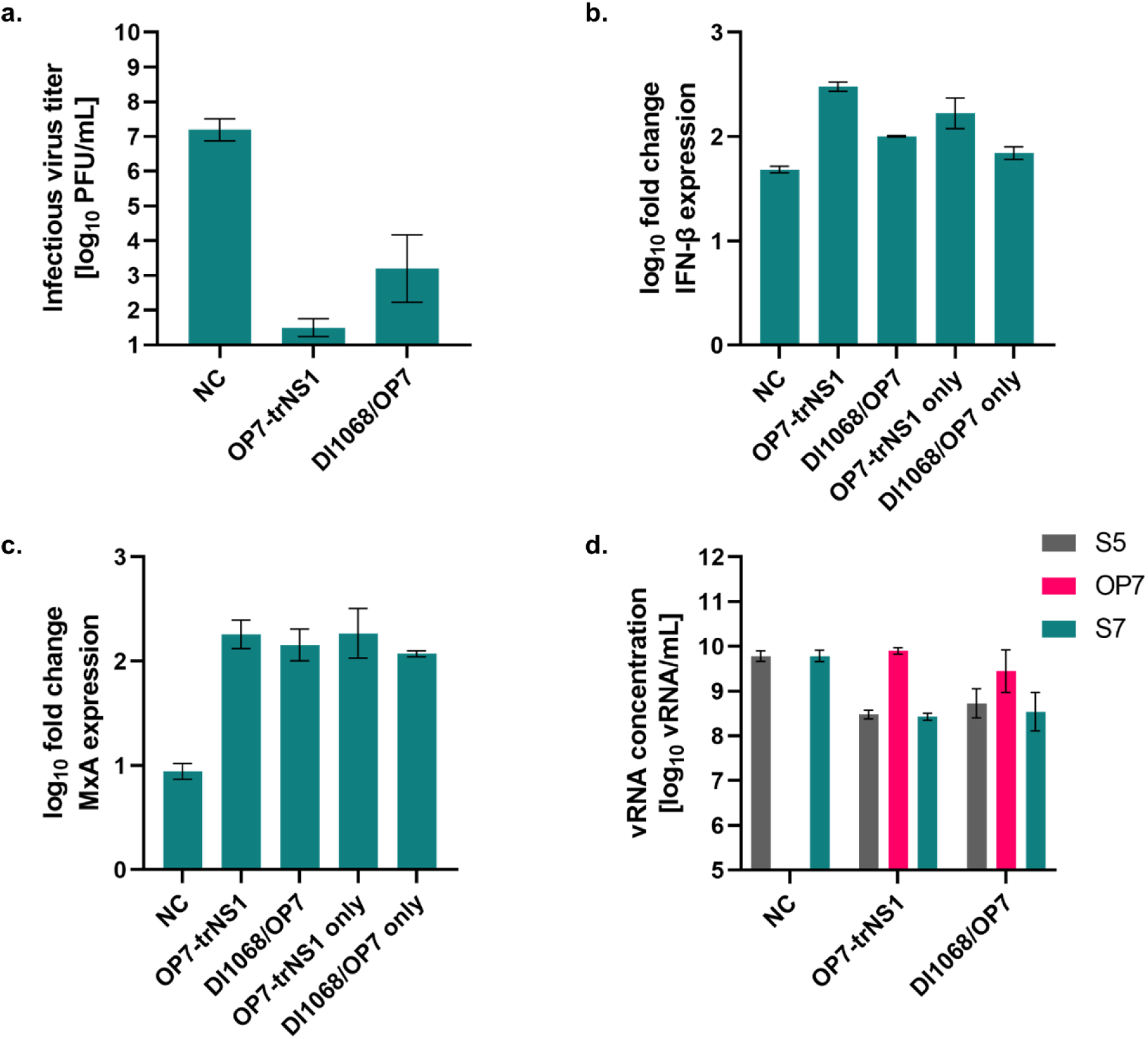
OP7-trNS1 enhances IFN induction significantly and increases antiviral activity by approx. two orders of magnitude. The produced and purified OP7-trNS1 material was subjected to an interference assay in Calu-3 cells. DI1068/OP7 was used as a control for comparison. Cells were infected with IAV PR8 alone at a MOI of 0.01 (NC) or coinfected with DIPs: OP7-trNS1 or DI1068/OP7. (a) Infectious virus release in the samples was measured by the plaque assay at 24 hpi. (b) IFN-β and (c) MxA expression at 24 hpi were investigated using RT-qPCR. (d) vRNA levels of progeny virions from cell culture supernatants were quantified by RT-qPCR. DIP input was normalized and calculated based on the HA titer. Error bars indicate the standard deviation of three independent experiments.

Overall, production MOIs influenced total virus yield, the fraction of OP7-trNS1 DIP and interfering efficacies. This data allowed cell culture-based production of OP7-trNS1 material with high antiviral activity.

### OP7-trNS1 exhibits enhanced antiviral activity and IFN induction in human lung cells

We next tested whether the OP7-trNS1 DIP preparation shows the predicted increase in antiviral activity and IFN induction in human lung epithelial cells. For this, the produced and purified OP7-trNS1 material was subjected to an interference assay to study and compare its antiviral activity to DI1068/OP7. Calu-3 cells were used for coinfection experiments to observe the effects in the presence of a functional IFN response to human IAV. Besides analysis of infectious virus titers, we quantified IFN-β and MxA expression using RT-qPCR analysis.

OP7-trNS1 coinfections showed a superior reduction of infectious virus release compared to DI1068/OP7 coinfections with IAV PR8 (Figure 7a). More specifically, OP7-trNS1 suppressed infectious virus titers over 100-fold more effectively than DI1068/OP7. Further analysis of gene expression in Calu-3 cells showed that OP7-trNS1 coinfections induced higher IFN-β levels compared to DI1068/OP7 coinfections (Figure 7b). Interestingly, IFN-β stimulation caused by OP7-trNS1 infection alone was also higher compared to DI1068/OP7 infection alone. This enhanced stimulatory effect was less pronounced at the level of MxA expression (Figure 7c). Results from vRNA quantification of progeny virions from cell culture supernatants showed an overproportional accumulation of S7-OP7 vRNA (Figure 7d). At the same time, the levels of S5 and S7 vRNAs were reduced, consistent with the phenotype of S7-OP7 vRNA-containing DIPs.

In summary, OP7-trNS1 reduced infectious IAV replication by approx. two orders of magnitude more effectively than DI1068/OP7 DIP and induced a stronger intracellular IFN response in Calu-3 cells. This enhanced antiviral activity was consistent with the model prediction and supports our conclusion that a NS1 truncation enhances DIP-mediated antiviral activity by reducing IFN antagonism.

## DISCUSSION

Herein, we used mechanistic modelling to guide the rational engineering of an IAV DIP with enhanced antiviral activity. A previously validated multiscale model predicted that removing NS1-mediated IFN antagonism would strengthen the antiviral effect of OP7. This prediction guided the design of OP7-trNS1, which was successfully reconstituted and produced in cell culture. In human lung epithelial cells, OP7-trNS1 induced a stronger IFN response and suppressed infectious IAV replication by approx. two orders of magnitude more effectively than the parental OP7 DIP, consistent with the model prediction. Together, these findings demonstrate that reducing viral immune antagonism is a viable strategy for rationally enhancing DIP antiviral activity.

Not all designed DIP genome constellations could be reconstituted. In this context, the introduction of large deletions and mutations into genome segments that do not naturally coexist in a single viral particle poses a risk of genetic incompatibilities [51]. The inability to assemble some genetic compositions is a known phenomenon observed in nature, as well as in the production of vaccine seed viruses using classical reassortment methods [52]. Here, most reassortment events lead to RNA-and protein-based incompatibilities between coinfecting viruses, which result in the production of progeny viruses with fitness defects, or in no reassortant virus [51]. For instance, suboptimal compatibility between the packaging signals of two parental viruses may limit, or even preclude, the packaging and assembly of new viruses [53]. From an RNA perspective, the formation of a new virus has been reported to be directed by vRNA structure [54]. This structure plays a role in the assembly of vRNP complexes (consisting of vRNA and viral protein), functional units packaged selectively into budding virions. Changes in the RNA sequence, such as those we introduced while designing novel DIP constructs (Figure 2), potentially affect the RNA-RNA interaction network, since some of the intersegment interactions drive vRNP co-segregation during assembly and budding [54]. This interaction network shows some level of plasticity, but can only accommodate a limited extent of sequence variation to assemble new gene constellations [54]. Due to the complexity of genetic interactions among the eight vRNPs, our understanding of genome assembly and its limitations remains incomplete. Therefore, outcomes are difficult to predict, which further underscores the challenges of novel DIP designs.

However, under these unfavorable conditions, we managed to balance the introduction of desired genetic alterations and the maintenance of virus fitness in the DI1068/NS124 and OP7-trNS1 constructs (Table 1, Supplementary Figure S1). Additional modifications to the reverse genetics procedure, such as introducing a point mutation in S7 vRNA (A14U) [46] and providing an NS1 protein expression plasmid (pCAGGS-NS1) [49], enabled the successful reconstitution of NS1-modified DIPs. With respect to the A14U substitution in S7 vRNA, previous studies showed that NS1 regulates S7 mRNA splicing (including that of M2 mRNA) and that IAV NS1 mutants were unable to express sufficient amounts of M2 mRNA [46]. The adaptive A14U substitution in S7 vRNA restores M2 splicing and expression in NS1 mutants and enhances their replication [46]. The addition of an NS1-expression plasmid (pCAGGS-NS1) during rescue was necessary due to its impaired expression in the NS1-modified DIPs. The absence of this plasmid resulted in no detectable virus rescues. Interestingly, NS1 expression *in trans* was required at the stage of virus reconstitution, but was not necessary at later production stages. During virus reconstitution, it appears that the IFN-competent HEK-PB2(adh) cells do not allow for replication of NS1-modified DIPs in the absence of NS1 due to enhanced antiviral IFN signaling. However, NS1-modified DIPs can grow in partially IFN-deficient MDCK-PB2(sus) cells during later production, regardless of the presence of NS1. This is likely due to the fact that MDCK cells are not able to exert an IFN-mediated antiviral response in this case, as the IFN-induced canine Mx1 protein cannot efficiently inhibit human IAV [49].

Nevertheless, cell culture production revealed that OP7-trNS1 grows to overall lower titers than DI1068/OP7 in MDCK-PB2(sus) cells, regardless of the conditions used. This observation is pointing to the growth attenuation previously observed among NS1 virus mutants. However, this attenuation was more prominent in cell lines capable of synthesizing IFN and conducting IFN signaling [37, 46–48, 55, 56], while in IFN-deficient Vero cells, 1 log reduction in the viral titers of the delNS1 virus compared to the WT virus was still reported [47]. In line with these observations, altered growth patterns were also observed for viruses containing truncated NS1 proteins in partially IFN-deficient MDCK cells [46–48], the cell line used in our reconstitution procedure. This can be explained by impaired accessory functions of the NS1 protein in the NS1 mutants. For instance, beyond its well-characterized IFN-antagonism, NS1 has also been implicated in the regulation of many processes in the viral replication cycle [33]. Although these additional functions are still poorly understood, the disruptions of accessory functions were frequently reported in NS1 mutant viruses [46, 48]. Nevertheless, in our studies, we obtained high production yields (HA titer of 2.62 log_10_ HAU/100µL), which are typically also observed in cell culture-based IAV production processes for use as vaccines [30, 57].

Modifications of the NS1 protein have an advantage over the full deletion of NS1 because viruses harboring them replicate to higher titers [48]. For instance, different NS1 truncations tested previously in pigs exhibited varying impacts on both NS1 function and expression. First, the length of the deletion determined which protein domains were retained and expressed, and, consequently, which protein functions were preserved or lost [55]. Second, the truncations also altered protein expression levels. Among truncated variants, the intermediate-length variant NS1Δ126, which is similar to the NS124 used in our studies, was reported to display the lowest level of NS1 expression [55]. This feature was considered beneficial, as lower expression diminishes the residual IFN antagonism while still providing the minimal NS1 levels required to support the protein functions essential for virus growth e.g. the aforementioned accessory functions. Together, these results suggest that domain-oriented removal critically influences virus replication and final titer. This was important for the high-yield production of the presented OP7-trNS1 DIP material (Figure 4).

The higher antiviral activity of OP7-trNS1 was consistent with gene expression analysis indicating higher IFN-β and MxA expression (Figure 7), and confirm the initial model predictions (Figure 1) of an enhanced IFN-induced antiviral state. Previously published reports on viruses with modified NS1 proteins have shown enhanced levels of type I IFN to be stimulated locally upon infection [37, 47]. This strategy was originally used to attenuate IAV for use as a LAIV [37]. In the present study, this feature was used to enhance the antiviral activity of our DIPs, as truncated NS1 variants shift the balance from immune evasion toward antiviral innate immune activation. This could facilitate the development of DIPs for use as a broad-spectrum antiviral against IFN-sensitive respiratory viruses. For instance, OP7 DIPs inhibited the replication of nonhomologous respiratory viruses, such as SARS-CoV-2 and RSV [27, 28, 44]. This inhibitory effect was mediated by the ability of DIPs to induce an enhanced IFN-mediated antiviral state. Similar findings have been reported by other research groups. Another IAV DIP, DI244, protected mice coinfected with the influenza B virus from severe disease, reducing both weight loss and clinical disease symptoms [29]. It also protected mice infected with a lethal dose of pneumonia virus from death [24]. A protective effect attributed to the stimulation of innate immunity has also been reported for non-IAV DIPs. Intranasal administration of poliovirus DIPs in mice elicited antiviral effects mediated by type I IFN responses against other enteroviruses, influenza viruses, and SARS-CoV-2 [25]. Broad-spectrum antiviral activity was also demonstrated for dengue virus DIPs against SARS-CoV-2, RSV, yellow fever virus, and Zika virus *in vitro* [26]. Again, the inhibition was shown to be mediated by IFN-dependent antiviral responses. Taken together, our findings suggest that further upregulation of IFN responses, by reducing IFN antagonism, might contribute to the development of DIPs as broad-spectrum antivirals that protect against both homologous and heterologous viral infections.

Beyond enhanced IFN production, NS1-modified viruses induce dendritic cell maturation [36] and stimulate T-cells [35]. In addition, they exhibit increased immunogenicity owing to the adjuvant properties of type I IFN [37]. As a result, non-DIP NS1 mutants have been widely explored as vaccine candidates, including LAIV [37]. They induced enhanced influenza virus-specific humoral and cellular immune responses in different animal models [55, 58–61], were well tolerated, and capable of inducing robust immune responses. Importantly, animals were not shedding the virus from the upper respiratory tract. The introduction of desired antigens into the current OP7-trNS1 construct may result in a highly promising live mucosal vaccine construct in the future. In general, mucosal administration offers several advantages over intramuscular vaccination. It induces broader and more durable mucosal and cell-mediated immune responses directly at the primary site of respiratory virus infection, apart from the systemic response [62–65]. A hypothetical live mucosal OP7-trNS1 vaccine may also provide advantages over existing LAIVs, which are administered intranasally but are based on attenuated rather than defective viruses. For these LAIV, residual virus replication may still cause disease [66–68]. In contrast, DIP-based live vaccines would have an improved safety profile, as they are replication-deficient, which could expand the range of vaccine recipients like elderly and immunocompromised individuals. In addition, previous studies have shown that inserting epitopes of non-influenza origin into NS1-mutant viruses induces robust immune responses in mice [69–73]. These findings indicate that the potential of DIPs harboring an NS1 truncation extend beyond the antiviral application presented in this study and that they could be developed towards use as mucosal vaccines.

Taken together, our findings establish OP7-trNS1 as a promising next-generation DIP with enhanced antiviral activity and IFN induction in human lung cells. Importantly, this study demonstrates that mechanistic systems virology can be used to rationally engineer DIPs by identifying and targeting specific viral determinants of antiviral activity, thereby complementing conventional trial-and-error approaches. The enhanced antiviral activity of OP7-trNS1 warrants further evaluation of its safety, prophylactic and therapeutic treatment windows and efficacy *in vivo.* Its increased IFN induction also provides a rationale for investigating its enhanced activity against other IFN-sensitive respiratory viruses. Together with the established cell culture-based production process, these findings provide a foundation for further preclinical development of OP7-trNS1. In addition, our study provides a proof-of-concept that mechanistic modelling can move DIP development from empirical optimization toward rational antiviral engineering.

## METHODS

### Mathematical multiscale model

A mathematical multiscale model, calibrated on intracellular and extracellular dynamics of STV and OP7 coinfection in Calu-3 cells, and able to consider antiviral effects upon IFN induction (described elsewhere [38]), was used to predict the impact of an OP7 unable to express NS1. To mimic an OP7-delNS1, we defined it as containing no functional S8 vRNA.

### Cells and viruses

MDCK cells (European Collection of Authenticated Cell Cultures (ECACC), #84121903) and MDCK-PB2(adh) cells (retrovirally transduced to express the IAV PB2 protein, as described previously [43]) were cultivated in Glasgow Minimum Essential Medium (GMEM, Thermo Fisher Scientific, #221000093) supplemented with 10% fetal bovine serum (FBS, Merck, #F7524) and 1% peptone (Thermo Fisher Scientific, #211709). For MDCK-PB2(adh) cells, puromycin (Thermo Fisher Scientific, #A1113803) was added as a selection marker to the medium at a final concentration of 1.5 μg/mL. HEK-293T-PB2(adh) cells (expressing PB2, as described previously [43]) were maintained in Dulbecco’s Modified Eagle Medium (DMEM) supplemented with 10% FBS, 1% penicillin/streptomycin (10,000 units/mL penicillin and 10,000 μg/mL streptomycin, Thermo Fisher Scientific, #15140122) and puromycin at a concentration of 1 μg/mL. Calu-3 cells (American Type Culture Collection (ATCC), #HTB-55) were cultured in DMEM/Nutrient Mixture F-12 (DMEM/F12) (Thermo Fisher Scientific, #11320033), supplemented with 10 % FBS, 1 % penicillin/streptomycin and 1 % non-essential amino acids (Thermo Fisher Scientific, #11140050). All adherent cells were maintained at 37°C and 5% CO_2_.

MDCK-PB2(sus) cells (retrovirally transduced to express PB2, as described previously [30]) were cultivated in chemically defined Xeno-CD001S MDCK CD Medium (Bioengine, #EXP0104103), supplemented with 0.5 μg/mL puromycin. Cultivation of the suspension cells was performed in shake flasks (125 mL baffled Erlenmeyer flask with vent cap, Corning, #1356244) in 50 mL of working volume in an orbital shaker (Multitron Pro, Infors HT; 50 mm shaking orbit) at 185 rpm, 37 °C and 5% CO_2_. To quantify VCC and viability, Vi-cell™ XR (Beckman Coulter, #731050) was used. IAV PR8 strain (provided by Robert Koch institute, #3138) was used as an infectious IAV strain in the interference assay. MOIs were determined based on the TCID_50_ titer (interference assay) and total virus particle concentration, derived from the HA titer (cell-culture based production of DIPs in shake flasks).

### Rescue of the DIP constructs

The generation of DIPs was based on a previously established plasmid-based reverse genetics system for the rescue of conventional PR8-derived DIPs harboring a deletion in S1 vRNA [43] and of OP7 chimera DIPs [7]. In order to complement for the missing expression of the PB2 protein, a co-culture of HEK-293T-PB2(adh) cells and MDCK-PB2(adh) cells was used for the plasmid transfections. For the rescue of OP7 chimera DIPs, a pHW-based plasmid harboring the sequence of S7-OP7 vRNA (GenBank accession number: MH085234) was used [7]. The pHW-based plasmids harboring the sequence of S1 vRNA DI1068, S7 vRNA with an A14U point mutation, S8 vRNA delNS1 and NS124 were generated in the present study. For this, the pHW2000 backbone plasmid [74] was kindly provided by Stefan Pöhlmann and Michael Winkler (German Primate Center, Goettingen, Germany). Different compositions of pHW-based plasmids were co-transfected depending on the designed construct, based on the modified procedure described previously [7]. In brief, plasmids encoding S1 DIP and DI1068 were transfected at 500 ng, S7-OP7 at 50 ng (always accompanied with S7 or S7 A14U in NS1 mutants at 1 μg), S8 delNS1 and NS124 at 1 μg, and the remaining WT pHW plasmids [74] at 1 µg using Lipofectamine 2000 (Thermo Fisher Scientific, #11668027) as a transfection agent according to manufacturer’s instructions. For the reconstitution of the NS1 mutants, an additional pCAGGS-based NS1 expressing plasmid (at 1 µg) [49] and S7 with A14U point mutation instead of WT S7 were used. After reconstitution, the rescued DIP material was amplified in MDCK-PB2(adh) cells.

### OP7-trNS1 and DI1068/OP7 DIP material production in shake flasks

Production of OP7-trNS1 and DI1068/OP7 DIP preparations in shake flasks using MDCK-PB2(sus) cells was performed with complete medium exchange prior to infection as described previously [7]. In brief, cells in the exponential growth phase were centrifuged (300×g, 5 min, RT) and resuspended in fresh medium (without puromycin) containing trypsin (final concentration 20 U/mL, Thermo Fisher Scientific, #27250-018) at a density of 2.0 × 10^6^ cells/mL. Cells were infected at indicated MOIs at 32 °C and 5% CO_2_. At indicated time points, samples were centrifuged (3000×g, 4 °C, 10 min) and supernatants were stored at-80 °C until further analysis. vRNA from progeny virions was extracted from these supernatants using the NucleoSpin RNA virus kit (Macherey-Nagel, #740956) according to the manufacturer’s instructions and stored at-80 °C until RT-qPCR, segment-specific RT-PCR, or sequencing.

The final OP7-trNS1 and DI1068/OP7 material tested in the interference assay in Calu-3 cells (Figure 7) was produced as described above at MOI of 1E-1 and 1E-2 (both harvested at 34 hpi), respectively. The produced DIPs were clarified (3000×g, 10 min and 4 °C), purified and concentrated by steric exclusion chromatography as previously described [31, 50]. Next, sorbitol (Roth, #6213) was added to the DIP material at a final concentration of 4%. Material was stored at − 80 °C until further use.

### Virus quantification

Total virus titers were quantified using the HA assay as previously described [75]. HA titers were expressed as log_10_ HA units per test volume (log_10_ hemagglutination units (HAU)/100 µL). The concentrations of DIPs were calculated using the HA titer and the equation below [76], where cRBC represents the red blood cell concentration (2.0 × 10⁷ cells/mL).

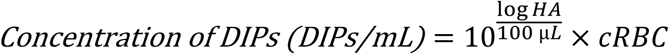

Infectious virus titers were quantified using the plaque assay and TCID_50_ assay using MDCK cells as previously described [30, 77].

### Segment-specific RT-PCR

To confirm IAV segment composition and to detect contaminating DI vRNAs in the DIP material (Figure 6), purified vRNA from progeny virions from cell culture supernatants was subjected to segment-specific PCR as described previously [10]. In brief, vRNA was subjected to reverse transcription using a universal “Uni12” primer [78] that binds to all eight genome segments. The resulting cDNA products were subjected to PCR using segment-specific primers to amplify each genomic segment individually. PCR products were analyzed by agarose gel electrophoresis.

### Quantification of vRNAs using RT-qPCR

In order to quantify the purified vRNAs from progeny virions from cell culture supernatants, we used a previously described RT-qPCR method that enables polarity-and gene-specific quantification of individual vRNAs [10, 79]. Tagged primers employed for the quantification of the vRNA of S5 are listed in [10], S7-OP7 in [31], and S7 WT in [7]. For the absolute quantification of vRNA concentrations, RNA reference standards were used, as described previously [10].

### Interference assay

The produced DIP material was tested for the interfering efficacy *in vitro* according to a previously established protocol [30, 41]. In brief, the inhibition of the infectious IAV PR8 virus propagation was investigated in a coinfection with DIP preparations. A MOI of 0.01 was used for the infectious virus challenge. DIP input was normalized based on the DIP concentration, calculated from the HA titer (described above). For the initial experiments with DI1068 and DI1068/NS124 (Figure 3), DIP material was used at a concentration of 1.4 E+09 particles per well. The material produced at different MOIs was used in the interference assay at a concentration of 1.02 E+09 particles per well for both OP7-trNS1 and DI1068/OP7 (Figure 5). In the final experiments with concentrated OP7-trNS1 and DI1068/OP7 (Figure 7), DIP preparations were applied at a concentration of 1.0 E+10 particles per well. After 24 hpi, supernatants were harvested and analyzed for infectious and total virus titers using the plaque and HA assay, respectively. RT-qPCR was used for the quantification of vRNA from progeny virions.

For the analysis of the IFN response, intracellular RNA was extracted. For this, the remaining cells were washed once with PBS, lysed with 350 μL RA1 buffer (Macherey-Nagel, #740955) and stored at-80 °C until RNA purification using the Nucleospin RNA isolation kit (Macherey-Nagel, #740955) according to the manufacturer’s instructions.

### Analysis of the IFN response using RT-qPCR

Expression of IFN-β and MxA of infected cells was assessed using RT-qPCR, as described previously [10, 41]. In brief, 500 ng of purified intracellular RNA was subjected to reverse transcription using an oligo(dT) primer and Maxima H Minus reverse transcriptase (Thermo Scientific, #EP0752) according to the manufacturer’s instructions. Subsequently, qPCR was performed using gene-specific primers for IFN-β and MxA [28] and 18S RNA as a reference housekeeping gene [41]. Gene expression was calculated using the ΔΔCT method [80] using the reference housekeeping gene 18S and expressed as fold change (relative to untreated, uninfected cells (mock infection control)).

## ACKNOWLEDGEMENTS

We thank Nancy Wynserski for excellent technical assistance.

## Author contributions

Conceptualization, J.P., D.R., U.R., S.Y.K.; Formal Analysis, J.P., D.R.; Funding Acquisition, U.R., S.Y.K.; Investigation, J.P., D.R.; Project Administration, J.P., S.Y.K.; Supervision, U.R, S.Y.K.; Visualization, J.P., D.R.; Writing – Original Draft, J.P.; Writing – Review & Editing, J.P., D.R., U.R., S.Y.K.

