## Supplementary information for "Model-Guided Engineering of NS1-Truncated Influenza OP7 Particles with Enhanced Interferon-Mediated Antiviral Activity"

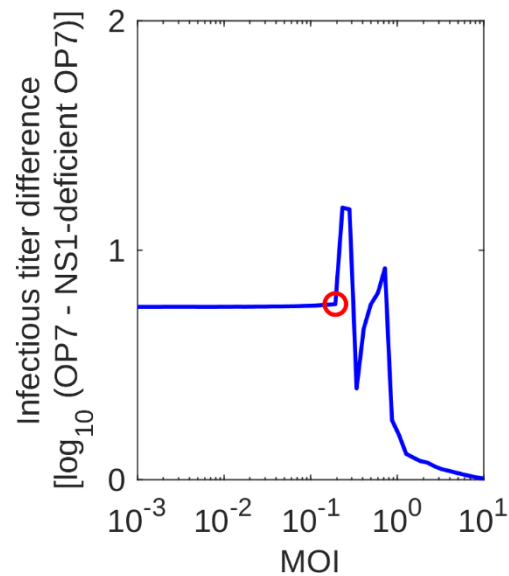

**Supplementary Figure S1. Comparison of OP7 and NS1-deficient OP7 inhibition at different STV MOI coinfections.**

Model simulations of STV infection and coinfections with OP7 or NS1-deficient OP7. Differences between infectious titers for coinfections with OP7 and NS1-deficient OP7 at varying STV MOIs. The difference for 36 hpi is shown. MOI 0.2 that was used to simulate dynamics in Figure 1B-C is shown as a red circle. Overall, lower MOIs lead to larger titer differences between OP7 and NS1-deficient OP7 coinfection than higher MOIs. Between MOIs of 0.23 to 1, the observed titer differences show sudden increases and decreases. This is caused by an artifact during model simulation, which would likely not be observable in a real infection scenario and is occurring due to our specific model setup.

The artifact can be explained by a specific design decision of the model. Due to the deterministic nature of the model, cells could theoretically release fractions of full virus particles, e.g., 1 % of an infectious virion. This could, then, already lead to the infection of new cells and would speed up the infection of the whole cell population drastically. This is not biologically accurate and, therefore, we assumed that the release of virus particles only starts when one full infectious progeny virion is formed and not when only a fraction of a particle was produced. This causes burst release events when all cells infected at the same time release their first virus particles leading to the step-like dynamics in Figure 1B. At low MOIs and MOIs above 1, two of these burst releases occur. At some MOIs between 0.3 and 1, three of these burst events occur causing a sudden jump in the infectious titer difference.

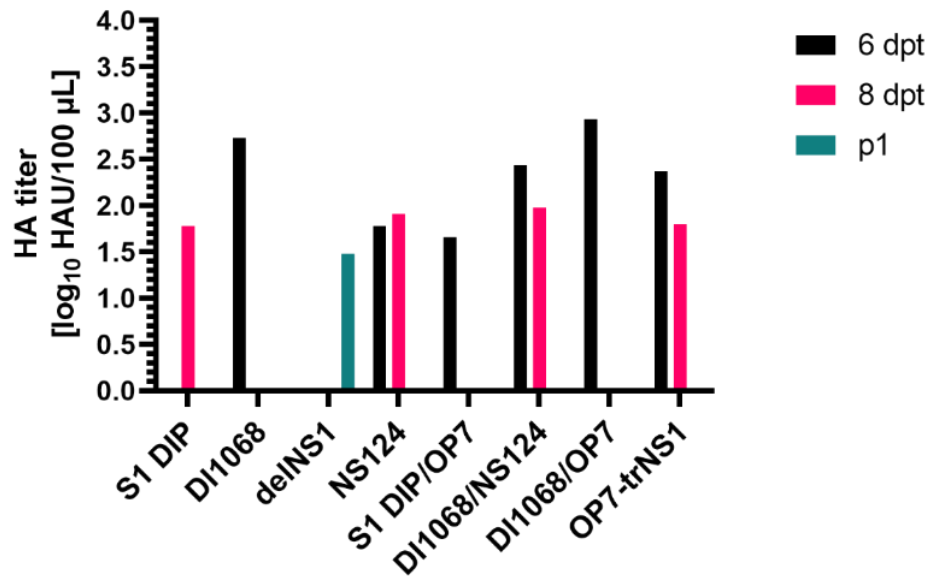

**Supplementary Figure S2. Total virus titers from virus reconstitution.**

HA titers were measured at 6 and 8 days posttransfection (dpt), and after the first round of subsequent virus passaging (p1) in MDCK-PB2(adh) cells. The time at which HA titers were detected for different constructs depended on the length of the deletion. The S1 DIP showed detectable HA titers 2 days later than DI1068 DIP carrying a smaller internal deletion in S1 vRNA. The delNS1 virus titer was detectable only after p1, indicating that in the presented reverse genetics system, a longer time is needed for the delNS1 virus propagation compared to NS124, which was detectable at 6 dpt. S1 DIP/OP7 and DI1068/OP7 showed detectable titers at 6 dpt. The newly generated complex constructs, DI1068/NS124 and DI1068/OP7/NS124 (OP7-trNS1), exhibited HA titers in the range between 2 and 3 log<sub>10</sub> HAU/100μL as early as 6 dpt. The early detection is consistent with the early rescue of respective mutants containing single and double genomic alterations. These mutants formed the basis for constructing the triple mutant OP7-trNS1. This early detection can be attributed to the reduced length of the S1 vRNA deletion and the point mutation that formed a stop codon instead of a deletion in the S8 vRNA sequence.
